# Transglutaminase mediated crosslinking of pea protein: from rheology to proteomics

**DOI:** 10.1101/2025.06.24.661300

**Authors:** A. Mukherjee, J. Van Pee, D. Duijsens, T. Grauwet, I. Van de Voorde, I. Fraeye, F. Weiland

## Abstract

Enzymatic crosslinking of pea proteins by transglutaminase (TG) is a promising processing strategy to enhance structure in protein-based systems. Despite its widespread application, detailed insights spanning macro- to molecular-level mechanisms remain limited. This study investigates TG-mediated crosslinking of pea protein by integrating rheological, chemical, and proteomic analyses to establish a comprehensive structure–function relationship. Dispersions of 10% w/v pea protein were treated sing TG dosages of 0–10 U/g protein. Rheological measurements showed a significant increase in storage modulus (G′) and altered gelation dynamics, with a clear dose-dependent relationship, indicating enhanced network formation. The content of free primary amino groups, quantified using the o-phthalaldehyde (OPA) method, decreased with higher TG levels, confirming progressive covalent crosslinking. SDS-PAGE further revealed differential participation of pea protein fractions in the crosslinking reaction. To elucidate molecular changes, proteomic profiling via LC-MS/MS was performed. The analysis indicated that glutamine and lysine residues located near glutamic acid or within disordered regions of peptides were preferentially crosslinked. Specific TG-catalyzed adducts were also identified and found to be enriched in disordered protein regions. Together, the findings demonstrate a clear mechanistic link between enzymatic crosslinking at the peptide level and functional alterations in network structure and viscoelastic properties. This integrative approach offers a molecular framework for the rational design of protein-based systems with tuneable mechanical properties, supporting innovation in both plant-based and conventional food applications.

## 1. Introduction

Proteins provide important techno-functional characteristics to many food products. In particular, protein gelation is a crucial functional property, of forming a 3D network which entraps water and other food constituents, giving integrity to the structure of the food product (Bryant & Julian McClements, 1998). One of the most widely used methods to induce protein gelation is thermal treatment (Tang et al., 2024). Animal-based proteins such as meat and egg proteins generally have excellent heat-induced gelation properties, which is partly attributed to the formation of disulfide bonds, resulting in the formation of a strong, covalently linked network (SMYTH et al., 1998; Felix et al., 2017; Hinderink et al., 2019). However, with the rising global population, there is an increasing interest, particularly in Western countries, to shift towards more plant-based dietary protein sources due to environmental and ethical reasons (Harwatt, 2019; van den Boom et al., 2023).

The main plant proteins currently used in commercial plant-based food products are soy and wheat gluten because of their fibrous structure and elastic texture (O’Sullivan et al., 2016; Snel et al., 2022). Pea protein is increasingly considered a promising alternative due to its hypo-allergenicity (in contrast to soy and wheat gluten), balanced amino acid profile, and broad availability (Boye et al., 2010). Pea seeds contain 19-25 % protein on wet basis, while globulins and albumins are the major storage proteins of which globulins account for 55-70 % (Boye et al., 2010; Freitas et al., 2000). Globulins are salt soluble proteins and consist of legumin (11S), vicilin (7S) and convicilin (7S) sub-classes. Legumins are hexameric proteins, with each monomer composed of a 40 kDa acidic and 20 kDa basic, (α-β) subunit linked by a disulfide bond. The quaternary structure is then connected with non-covalent bonds. On the other hand, vicilins are trimers with subunits of 55, 50, 35 and 17 kDa. Lastly, convicilin is a trimeric protein with a molecular mass of 71 kDa for each monomer. The other major storage protein fraction in pea is water-soluble albumins accounting for 10-20 % of the total storage proteins (Duranti & Scarafoni, 1999; Higgins et al., 1986). The two main albumins are pea albumin 1 (PA1) and pea albumin 2 (PA2), with molecular masses of around 6 kDa and 25 kDa, respectively. PA1 exists as a monomer, whereas the subunits of PA2 are possibly joined by non-covalent interactions to form a dimeric structure (Gruen et al., 1987). Although the inclusion of pea protein in plant-based foods is gaining interest, it is still not widely used commercially. This is in part due to its poor structure-forming capacity compared to other plant proteins such as soy (Xiong et al., 2018). While soy protein forms strong, covalent disulfide bridges upon heating, the pea protein network structure is established by comparatively weaker hydrophobic interactions and hydrogen bonds (Mession et al., 2015; Sun & Arntfield, 2012; O’Kane et al., 2004).

Previous studies have attempted to improve the pea protein network structure by the utilization of transglutaminase (TG, E.C. 2.3.2.13)(Sun & Arntfield, 2011). TG is an acyl transferase, facilitating the formation of covalent isopeptide bonds by linking γ-carboxamide groups of glutamines (acyl donor) with free amino groups (acyl acceptor). Specifically, ε-amino groups of lysine residues serve as acyl acceptors, resulting in the formation of intra- and intermolecular ε-(γ-glutamyl)-lysine cross-links (Djoullah et al., 2018; Duarte et al., 2020; Schlangen et al., 2023). Djoullah et al., (2015a), (2018) explored the effect of TG crosslinking of native and denatured pea protein sub-fractions. Liu et al., (2024) reported that TG treatment of cold-set pea protein gels after pH 12-shifting and heat treatment could improve the gelation property of the protein. Sun et al., (2011) reported formation of stable gels using TG treatment of salt extracted pea protein. Yaputri et al., (2024) compared gel properties of TG treated pea and chickpea protein under selected conditions of 3.3 nkat TG /mL, for 5 min. They found that the crosslinked chickpea had a significantly higher gel strength but a lower emulsification capacity than the pea protein. Nivala et al., 2017 reported that faba bean was crosslinked extensively with 10 nkat/g TG, whereas oat protein was crosslinked with 100-1000 nkat/g TG. Although these prior works have attempted to provide insight into the application of transglutaminase in crosslinking of pea protein, there is a lack of in-depth rheological data to characterize the evolution of gel formation and the quality of the gels formed, e.g., using frequency sweep and the determination of the linear viscoelastic region (LVR) of the gels. Moreover, the previous studies focused on macroscopic properties of the gel systems with little emphasis on the underlying molecular mechanisms. While mass spectrometry-based proteomics can be applied to study protein modifications, only limited attempts to study crosslinked proteins using this technique have been conducted due to the associated challenges of solubilization and hydrolysis of crosslinked proteins, and the complexity of data analysis (McKerchar et al., 2019; Petrotchenko et al, 2010). Thus, an opportunity exists to use MS-based proteomics to explore the effects of TG-mediated crosslinks at a molecular level to determine the localization of crosslink sites. In particular, proteome analysis can be used to determine the localization of crosslinks and to reveal whether certain structural domains, such as flexible disordered regions or certain amino acid motifs (short sequence–structure pattern), are preferentially modified by TG crosslinking, as was explored in this study.

In addition, importantly, commercial extraction procedures applied to isolate pea protein have a significant negative influence on the pea protein functionality, especially solubility and gelation properties, by imparting denaturation, resulting in protein aggregation (C. Kornet et al., 2020; Tanger et al., 2020; J. Yang et al., 2021). Recent studies emphasize the need for mild extraction procedures to produce soluble, native protein with high functionality (R. Kornet et al., 2021). Pea protein produced by membrane filtration such as ultrafiltration has been reported to have higher solubility compared to isoelectric precipitation (Boye et al., 2010; Timmermanns et al., 1993; Vose, 1980) Since the solubility of the protein affects the accessibility of lysine and glutamine to TG, it thus impacts the enzyme’s ability to catalyze crosslinking reactions effectively.

A detailed study to elucidate pea protein crosslinking by TG, combined with the identification of specific protein targets, is crucial to optimizing its application in food systems. The overarching aim of this study is to unravel the effect of different TG dosages on the structure-forming property of highly soluble, ultrafiltered pea protein, integrating traditional chemical and rheological analyses with advanced proteomics analyses.

## 2. Materials and methods

### 2.1. Materials

Dehulled yellow peas were received from Braet - De Vos NV (The Netherlands). A Ca^2+^-independent TG enzyme from *Streptomyces mobaraensis* was obtained from Novozymes (Galaya® Prime, Denmark). The enzyme exhibited an activity of 105.8 U/g as determined by the method of Djoullah et al. (2015a). All chemicals used were of analytical grade (unless mentioned otherwise).

### 2.2. Pea protein extraction

Pea protein extraction was performed using alkaline solubilization followed by purification using ultrafiltration and diafiltration. Dehulled yellow peas were milled using a rotor mill (Retsch Ultra Centrifugal Mill ZM200, Haan, Germany) with a 500 µm sieve. The resulting pea flour was dispersed in tap water at a 1:5 ratio (w/v) and adjusted to pH 9.5 using 6 M NaOH (Chem-Lab, Belgium). The dispersion was stirred for 1 h. Subsequently, the protein dispersion was centrifuged (Sorvall Lynx 6000, Thermo Scientific, Waltham, MA, USA) at 4500 g for 20 minutes at room temperature (RT). The supernatant was ultrafiltered (Alfa Laval TesUnit 20M, Lund, Sweden) using a membrane with a 25 kDa cut-off and a volume concentration factor of 5. The retentate was then subjected to diafiltration twice for further protein purification. The resulting filtrate was neutralized using 6 M HCl (Sigma-Aldrich, St. Louis, MO, USA) and lyophilized (Epsilon 2-10D LSCplus, Christ Systems, Osterode am Harz, Germany) to obtain the purified pea protein extract. Freeze-dried extracts from different extraction batches were then pooled into a single homogenous batch. The lyophilized pea protein extract was vacuum-packed into several aliquots and stored at -20 °C until further analysis.

### 2.3. Pea protein extract characterization

#### 2.3.1. Proximate composition

The protein content of the obtained pea protein extract was determined using the Kjeldahl method (ISO 20483) with a nitrogen to protein conversion factor (c.f.) of 5.44 (Mariotti et al., 2008). The total lipid content of the extract was measured gravimetrically after extraction using chloroform/methanol (1:1) (Ryckebosch et al., 2012). Oven drying was used to determine the moisture content of the protein extract (ISO 24557), and total mineral content was determined using dry ashing (ISO 2022). All analyses were conducted in triplicate.

#### 2.3.2. Differential Scanning calorimetry

The nativity of the protein in the extract was studied using a Q2000 heat flux differential scanning calorimeter with Advanced Tzero™ technology (TA Instruments, Delaware, USA) equipped with a refrigerated cooling system (RCS 90). The method was based on previous work by Chigwedere et al., 2018. Denaturation enthalpy, onset temperature and denaturation temperature were determined from the obtained thermograms using the TA universal analysis software 2000 (TA Delaware, USA). The analysis was conducted in duplicate.

#### 2.3.3 Solubility

Protein solubility was determined in triplicate according to Tanger et al., 2020, with modifications. The pea protein extract was dispersed in a 50 mM sodium phosphate buffer containing 0.3 M NaCl, pH 7 in a concentration of 1 % (w/v), and stirred at 500 rpm for 1 hour at RT using a magnetic stirrer. The dispersion was then centrifuged at 8960 g for 20 minutes at 20 °C (Hettich 1406, Tuttingen, Germany). The supernatant was collected, and its protein content was analyzed using the Kjeldahl method (section 2.3.1). Protein solubility was expressed as follows:

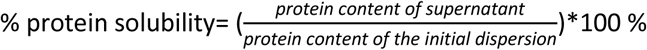

### 2.4. Enzymatic crosslinking of pea protein with TG

Pea protein extract was dispersed in phosphate buffer (50 mM sodium phosphate containing 0.3 M NaCl, pH 7), to a concentration of 10 % protein (w/v) and stirred at 500 rpm for 1 hour using a magnetic stirrer. An aliquot of 2 mL of the dispersion was transferred into reaction tubes, and TG was added in dosages of 0.25, 0.5, 1, 5, and 10 U TG/g protein. The samples were then incubated in a shaking water bath at 40 °C for up to 5 hours. At pre-set time points, a single individual tube was transferred to a water bath and incubated at 70 °C for 10 minutes to inactivate the enzyme. For blank samples, i) no enzyme was added to the protein dispersion, but the sample was heated at the inactivation conditions (hereafter referred to as “heat and no enzyme” (HNE)) ii) no enzyme was added to the protein dispersion and no heating step was conducted (hereafter referred to as “no heat and no enzyme” (NHNE)). The samples were subsequently freeze-dried and stored at -20 °C until further analysis.

### 2.5. Rheological analyses

Structure formation due to TG crosslinking of pea protein extract was analysed by measuring viscoelastic properties through small amplitude oscillatory shear rheology using a stress-controlled rheometer (AR 2000ex, TA instruments, New Castle, USA) on cross-hatched parallel plate geometry (40 mm diameter). Protein dispersions were prepared, and different dosages of TG were added as described in section 2.4, including a blank (0 U TG/g protein). The resulting dispersions were briefly vortexed and immediately loaded on the rheometer at 20 °C. For all analyses, the G’ (storage modulus), G” (loss modulus) and tan δ (G”/G’) were monitored. All analyses were conducted in duplicate.

#### 2.5.1. Time sweep

After sample application, the temperature of the plates was ramped up to 40 °C (5 °C/min). After this, a time sweep measurement was conducted for 5 h to detect structure formation as a function of incubation time. Then, the temperature was increased to 70 °C at 5°C/min and held for 10 min to inactivate the enzyme. Following the inactivation, the temperature was reduced to 20 °C at 5 °C/min and held for 30 min. Constant strain value of 0.5 % and frequency of 1 rad/sec were maintained during the entire time sweep. Particular time points were selected, namely end of incubation, end of inactivation, and process end for further comparison of G’ values.

#### 2.5.2. Frequency sweep

Frequency sweep measurements were performed on the gels formed immediately after the time sweep in a frequency range of 0.062 to 62 rad/sec with 20 measuring points per decade. The strain was kept constant at 0.5 % and the temperature at 20 °C.

#### 2.5.3. Strain sweep

Lastly, the samples underwent a strain sweep to determine the linear viscoelastic region (LVR) of the gels. The samples were subjected to strain values ranging from 0.01 to 1000 % at a constant temperature of 20 °C and a frequency of 1 rad/sec with 20 measuring points per decade. The extent of the LVR was determined by calculating the oscillatory strain at which the corresponding G’ deviated by more than 10 % from the average G’ of the preceding five measurements.

#### 2.5.4. Data analysis

The results from the frequency sweep were modelled using the Power law model (Ikeda et al., 1999):

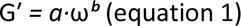

Where *a* is the consistency factor representing gel strength (Pa·s^b^), and *b* is the exponent indicating the dependence of G′ on frequency (ω, rad/s).

All values were estimated in duplicate. Furthermore, the difference between means of the rheological parameters during time sweep and strain sweep was tested using a one-way ANOVA, followed by an all-pair Tukey’s HSD test at 5 % significance level. The analysis was done using JMP Pro 17 (SAS Institute Inc. Cary, NC, USA).

### 2.6. Determination of free primary amino groups

#### 2.6.1. Sample preparation and spectrophotometric analysis

The free primary amino groups in the samples were quantified using o-phthaldialdehyde (OPA) (Djoullah et al., 2015b; Nielsen et al., 2001). The NHNE, HNE, and the crosslinked protein sample (section 2.4) (25 mg) were dispersed in 500 µl of 0.1 M sodium phosphate buffer containing 2 % (w/v) sodium dodecyl sulphate (SDS) (pH 8) and vortexed for 30 sec. The dispersion was centrifuged at 5641 g for 10 minutes at 20 °C (Centrifuge 5424 R, Eppendorf, Germany). The supernatant was diluted 1:20 (v/v) using 0.1 M sodium phosphate buffer containing 2 % (w/v) SDS (pH 8) and vortexed. Subsequently, 3 mL of freshly prepared OPA reagent (Nielsen et al., 2001) was mixed with 400 µL of the diluted supernatant, followed by an incubation for 2 min at RT in the dark. Afterwards, the absorbance was measured at 340 nm using a spectrophotometer (Cary 100 UV-Vis, Agilent Technologies, Santa Clara, CA, USA). An L-serine standard curve (25–150 mg/mL) was used to convert absorbance into L-serine equivalents and the concentration of free amino groups.

#### 2.6.2. Data analysis

Kinetic modelling of the decrease in free primary amino groups was done using SAS version 9.4 (SAS Institute, Inc., Cary, NC, USA). The data were modelled using a fractional conversion model (Verkempinck et al., 2024) as follows:

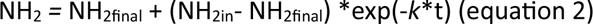

where, NH_2final_ is the final content of primary amino groups (mg/g sample), NH_2in_ is the content of primary amino groups (mg/g sample), *k* is the estimated rate constant of crosslinking (min^-1^), and t is the incubation time (min). Since each time point in the crosslinking study was assessed using an individual tube, the measurements can be considered independent observations of the system at distinct time points, enabling kinetic modelling of the data (Duijsens et al., 2022).

### 2.7. SDS-PAGE

The NHNE, HNE samples, as well as crosslinked samples (section 2.4), were dispersed in phosphate buffer (50 mM sodium phosphate containing 0.3 M NaCl, pH 7) at a concentration of 5 mg/mL and vortexed for 1 min. Protein dispersions were then mixed in a 1:1 ratio with sample buffer, containing 95 % Laemmli Sample buffer (Bio-Rad, Hercules, CA, USA) and 5 % β-mercaptoethanol (Bio-Rad, Hercules, CA, USA), incubated at 95 °C for 5 min, and afterwards cooled to RT. Samples as well as protein standards (Precision Plus Protein Unstained, Bio-Rad, Hercules, CA, USA) were loaded on the gel (Bio-Rad, Criterion TGX, Any KD^TM^, Stain-Free, Hercules, CA, USA). The electrophoresis chamber was filled with 800 mL Tris/Glycine/SDS buffer (Bio-Rad, Hercules, CA, USA). Electrophoresis was conducted at a constant current of 20 mA for 25 minutes, followed by a constant voltage of 300 V for 20 minutes. Finally, the gels were scanned using the ChemiDoc MP Imaging System (Bio-Rad, Hercules, CA, USA) using the stain-free settings.

### 2.8. Proteome analysis

#### 2.8.1. Hydrolysis of crosslinked proteins

##### Unfractionated cross-linked protein sample

The lyophilized crosslinked protein sample (section 2.4) (incubated with 10 U TG/g protein for 300 min, in quadruplicate) and the NHNE sample (in quadruplicate) were dispersed (10 mg/ml) in a buffer containing 8 M urea (Merck KgAA, Darmstadt, Germany) and 50 mM ammonium bicarbonate (Sigma-Aldrich) (pH 8.5) and solubilized using an ultrasonication rod (2 pulses of 30 sec each, 50 % amplitude, 1 min, 120 W) (Thermo Scientific, Waltham, MA, USA). The protein concentration of the solution was determined using a Bicinchoninic Acid (BCA) assay (Pierce™ BCA protein assay kit, Thermo Scientific Waltham, MA, USA). 50 µg of the proteins were reduced and alkylated for 45 min at RT using 25 mM chloroacetamide (Sigma-Aldrich) and 10 mM Tris(2-carboxyethylphosphine) (Sigma-Aldrich). One µg of Trypsin/LysC (Thermo Fisher Scientific, Waltham, MA, USA; Cat. no. A40007) was added to the proteins and incubated overnight at 37 °C. Finally, the samples were cleaned up using in-house packed C18 tip columns (Empore SPE Disks, 3M, Saint Paul, MN, USA), and freeze-dried.

##### High molecular weight (HMW) prefractionated crosslinked sample

Lyophilized crosslinked samples treated with either 0.25, 0.5, 1, 5 or 10 U TG/ g protein (section 2.4) were solubilized, and the protein content was determined as above. Then, 50 µg of the protein was loaded on SDS-PAGE gels under reducing conditions as described in section 2.7. Afterwards, proteins were fixed overnight in 40 % (v/v) ethanol (Chem-Lab Analytical, Zedelgem, Belgium, Cat. No. CL00.1807.5000) and 10 % (v/v) acetic acid (VWR, West Chester, PA, USA, Cat. No. 20104.298). Coomassie blue G250 (Bio-Rad laboratories, Hercules, CA, USA, Cat. No. 1610786) was used to visualize the proteins. High molecular weight gel bands were excised using a scalpel, cut into pieces of approximately 1 mm^3^, incubated in 30 % (v/v) acetonitrile (Merck, Cat. No. 1.00029.1000), 25 mM ammonium bicarbonate (Sigma-Aldrich), and destained for 30 min in an ultrasonic bath (Sonorex Digitec DT 52, Bandelin, Berlin, Germany). This step was repeated twice or until the gel bands appeared clear. Subsequently, the gel pieces were dehydrated using 100 % acetonitrile (Merck). The acetonitrile was then removed, and the proteins were reduced and alkylated as mentioned above. After reduction and alkylation, the liquid supernatant was removed, and the gels were dehydrated using 100 % acetonitrile (Merck). One µg of Trypsin/LysC (Thermo Scientific, Waltham, MA, USA) was added to the gel pieces and incubated overnight at 37 °C. Subsequently, the peptides were extracted using ultrasonication after sequential addition of 100 µl 1 % (v/v) formic acid (Merck), 100 µl of 60 % (v/v) acetonitrile (Merck), 1 % (v/v) formic acid (Merck), and finally 150 µl 100 % acetonitrile (Merck). The supernatants were collected, combined, and freeze-dried.

#### 2.8.2. TMT labelling

Isobaric labelling of the hydrolyzed peptides was conducted using Tandem Mass Tag (TMT) Pro 16-plex reagents (Thermo Scientific, Waltham, MA, USA). In brief, 0.5 mg of the TMT reagents were dissolved in 32 µl of anhydrous acetonitrile (Sigma-Aldrich) each and reconstituted according to manufacturer’s protocol. Freeze-dried peptides were resuspended in 50 mM 2-[4-(2-hydroxyethyl) piperazin-1-yl] ethanesulfonic acid (HEPES) (Sigma Aldrich) pH 8.5 (Zecha et al., 2019) and 6 µl of the respective TMT label was added. The labelling reaction was conducted for 2 h at RT and quenched via the addition of hydroxylamine (Sigma Aldrich) to a concentration of 0.5 % (w/v). The labelled peptides were then combined, freeze-dried, and finally resuspended in 2 % (v/v) acetonitrile (Merck) and 0.1 % (v/v) trifluoroacetic acid (TFA, Merck).

#### 2.8.3. Liquid chromatography -tandem mass spectrometry (LC-MS/MS) analysis

Peptides were re-constituted in 20 µl loading solvent A (0.1% (v/v) TFA in water/acetonitrile (ACN) (98:2, v/v)) for further LC-MS/MS analysis (6 µl injection volume for the unfractionated and 2 µl for the HMW prefractionated sample). LC analysis was performed using an Ultimate 3000 RSLCnano system (Thermo Scientific, Waltham, MA, USA). Peptides were trapped at 20 μl/min for 2 min in loading solvent A on a 5 mm trapping column (Thermo scientific, Waltham, MA, USA, 300 μm internal diameter (I.D.), 5 μm beads). The peptides were separated on a 250 mm Aurora Ultimate, 1.7µm C18, 75 µm inner diameter (Ionopticks) kept at a constant temperature of 45°C. Peptides were eluted by a non-linear gradient starting at 0.5% MS solvent B reaching 33% MS solvent B (0.1% FA in water/acetonitrile (2:8, v/v)) in 135 min, 55% MS solvent B (0.1% FA in water/acetonitrile (2:8, v/v)) in 155 min, 70% MS solvent B in 160 minutes followed by a 5-minute wash at 70% MS solvent B and re-equilibration with MS solvent A (0.1% FA in water). The LC system was in-line connected to a mass spectrometer (MS) (Q Exactive HF (Thermo Scientific, Waltham, MA, USA)) which was operated in data-dependent mode with MS/MS acquisition for the 12 most abundant ion peaks per MS full-scan spectrum. Full-scan MS spectra (375-1500 m/z) were acquired at a resolution of 60,000 in the Orbitrap analyzer with an ion accumulation AGC target value of 3E6. The 12 most intense m/z features above a threshold value of 1.3E4 were isolated (isolation window of 1.5 m/z) for fragmentation at a normalized collision energy of 33% (AGC target value of 1E5 for maximum 80 ms). MS/MS spectra (fixed first mass at 120 m/z) were acquired at a resolution of 45,000 in the Orbitrap analyzer. Polydimethylcyclosiloxane background ion (445.120028 Da) was used for internal calibration while QCloud (Chiva et al., 2018; Olivella et al., 2021) was used for the control of instrument longitudinal performance.

#### 2.8.4. Data analysis – Unfractionated cross-linked protein sample

The resulting spectra were searched against a *Pisum sativum* protein database (downloaded from www.uniprot.org on 02/10/2023; 64,176 entries) using Comet (version 2022.01 rev. 0) (Eng et al., 2015) and the Trans-Proteomic Pipeline (version 6.3.0 Arcus) (Panchal et al., 2024). For Comet analysis, the default parameters were used. In brief, precursor mass tolerance was set to 20 ppm, the enzyme specified as Trypsin, omitting the proline rule (Rodriguez et al., 2008), with a maximum of 2 missed cleavages. As variable modification, the oxidation of methionine was set. As fixed modifications, TMTpro labelling of the peptide N-terminus and lysine were specified. PeptideProphet was run using its default settings, with additional parameters including accurate mass, expectation score, and the specification of decoy hits. For the subsequent iProphet (Shteynberg et al., 2011) analysis, default settings were applied. For peptide/protein quantification, the raw file was converted to a centroided mzML file using ProteoWizard (Chambers et al., 2012) from which the TMT reporter intensities were extracted, and subsequently transformed and calibrated to each other using variance stabilizing normalization (Huber et al., 2002). Linear models for microarrays (limma) were employed to determine between-sample differences in the mean abundance of the crosslinked peptides (Ritchie et al., 2015), employing a 5 % false discovery rate (FDR) cut-off. Sequence motifs were identified using rmotifx (Wagih et al., 2016), while enrichment of disordered regions was investigated using a Fisher’s exact test. All data analysis was conducted using in-house written scripts (or modified from Jadav et al., 2024.) using R (version 4.3.1) in the RStudio environment (2024.12.1 Build 563). The following R libraries were used: stringr (Wickham, 2023), MSnbase (Gatto et al., 2012; Gatto et al., 2021), dplyr (Wickham, et al., 2023), ggplot2, reshape2 (Wickham, 2007), vsn (Huber et al., 2002), limma (Ritchie et al., 2015), seqinr (Charif et al., 2007), plyr (Wickham, 2011), ggrepel (Slowikowski, 2024), ggpointdensity (Kremer, 2019), scales (Wickham, Pedersen, et al., 2023), wesanderson Ram et al., 2023), matrixStats (Bengtsson, 2024), ggseqlogo (Wagih, 2024), and viridisLite (Garnier et al., 2023). All raw data and analysis files are available at ProteomeXchange (Deutsch et al., 2023) and jPOSTrepo (Okuda et al., 2017) with the identifiers PXD064878 and JPST003865, respectively. Data analysis scripts can be downloaded from Zenodo under https://doi.org/10.5281/zenodo.15640883.

#### 2.8.5. Data analysis – High molecular weight prefractionated cross-linked protein sample

MS raw data were searched using pLink2 (Chen et al., 2019) against the same FASTA as described above, with the addition of common contaminants (as per the FragPipe search engine). As a cross-linker, transglutaminase was specified (loss of NH_3_). Fixed modification was set as the carbamidomethylation of cysteine and TMTpro labelling of the peptide N-terminus. As variable modifications, TMTpro labelling of lysine and oxidation of methionine was set. Further, the MS raw data was searched using FragPipe (version 22) to QR and QK remnants on lysine residues. The same FASTA as used for the pLink2 search, with addition of reverse sequences, was used. Default FragPipe settings were applied, in brief, precursor and fragment mass tolerance were set to +/- 20 ppm. “Strict trypsin” was set as enzyme, omitting the proline rule (Rodriguez et al., 2008). As variable modifications, oxidation of methionine, protein N-terminal acetylation, and protein/peptide N-terminal TMTpro labelling were set. Further variable modifications of lysine were set: TMTpro, addition of a TG cross-linked QR (glutamine-arginine remnant) and QK (glutamine-lysine remnant). A cross-linked lysine remnant was set as variable modification of glutamine. Fixed modification was specified as carbamidomethylation of cysteine. The resulting peptide identifications were re-scored with MSBooster (K. L. Yang et al., 2023), using the DIA-NN retention time (version 1.8.2 beta 8) (Yu et al., 2023) and Prosit 2020 Intensity TMT deep learning models (Gabriel et al., 2022). For false discovery rate calculation, the Percolator algorithm (Käll et al., 2007; The et al., 2016) was employed, set to 1 % for peptide-to-spectrum matches (PSM). ProteinProphet (version 5.1.1) (Nesvizhskii et al., 2003) was applied for protein identification. Results (PSM, peptides, proteins) were filtered using Philosopher with a 1 % FDR (da Veiga Leprevost et al., 2020). The MS raw data was also searched using FragPipe (version 22) (Kong et al., 2017) to identify QR and QK remnants on lysine residues-link sites ProteomeXchange and jPOSTrepo under the identifiers PXD064875 and JPST003864, respectively. All analysis scripts are available via Zenodo under https://doi.org/10.5281/zenodo.15640016. All MS raw data and search engine results are available via ProteomeXchange and jPOSTrepo under the identifiers PXD064875 and JPST003864, respectively. Data was analysed using R (version 4.3.1) in the RStudio environment (2024.12.1 Build 563) using in-house scripts written as described above.

## 3. Results and discussion

### 3.1. Characterization of ultrafiltered pea protein extract

The protein content of the ultrafiltered pea protein extract was determined to be 74.1 ± 1.0 %. The protein content obtained is slightly higher than previously reported values for pea protein obtained by ultrafiltration by Mondor et al., 2012 (72.1 %, recalculated using 5.44 c.f. and as the average from 3 ultrafiltration modules) but lower than that reported in Taherian et al., 2011 (82.2%, recalculated with 5.44 c.f.). The lipid, ash, and moisture content of the extract were measured to be 10.259 ± 0.004 %, 5.99 ± 0.04 %, and 2.4 ± 0.3 %, respectively. The denaturation enthalpy of the pea protein extract was observed to be 10.1 ± 0.3 J/g protein with a denaturation temperature of 87.06 ± 1.01 °C. The onset of denaturation occurred at 77.4 ± 0.5 °C. It is difficult to compare values of denaturation temperature across the literature since denaturation is a time-temperature effect. Hence, it depends on the experimental conditions under which the parameters were measured (Ricci et al., 2018). It is also known that the protein extraction method impacts the resulting denaturation enthalpy (Tanger et al., 2020; Verkempinck et al., 2024). The denaturation enthalpy values obtained in this study are higher than those reported for alkali extractions (Shand et al., 2007; Tanger et al., 2020), but lower than those for salt-extracted pea protein (Sun & Arntfield, 2010). However, the denaturation enthalpies obtained in this study suggest that the nativity of the protein during the extraction procedure was largely retained.

The ultrafiltered protein extract exhibited a solubility of 90.79 ± 1.02 %, which is in line with retention of nativity in the ultrafiltered extract, as solubility values of above 80% are obtained when applying mild protein extraction methods such as salt extraction and micellar precipitation (Stone et al., 2015).

### 3.2. Rheological analysis

Gel formation was assessed via gelation kinetics and viscoelastic properties of pea protein extract (10% w/v) incubated with varying TG dosage (0 – 10 U TG/g protein) using small amplitude oscillatory shear rheology. **Fig. 1 A** represents the evolution of the storage modulus (G’) during the incubation of the protein dispersions with TG at 40 °C for 5 h, subsequent thermal enzyme inactivation step at 70 °C for 10 min, and cooling. For all the systems, G’ was higher than G” (the loss modulus) (results for G” are not shown), indicating gel formation. Djoullah et al., (2018) reported similar trends of G’ evolution for pea protein dispersions treated with TG. However, they did not explore the effects of different TG dosages, thermal inactivation, and subsequent cooling on the G’ of the protein systems. The G’ of all the samples, including that of the blank, rose further during the inactivation step, owing to the additional network development, i.e., thermal gelation within the protein systems due to the input of thermal energy (Kornet et al., 2021; Sun et al., 2012). Finally, G’ values stabilized at the end of the process after cooling down.

**Figure 1.**
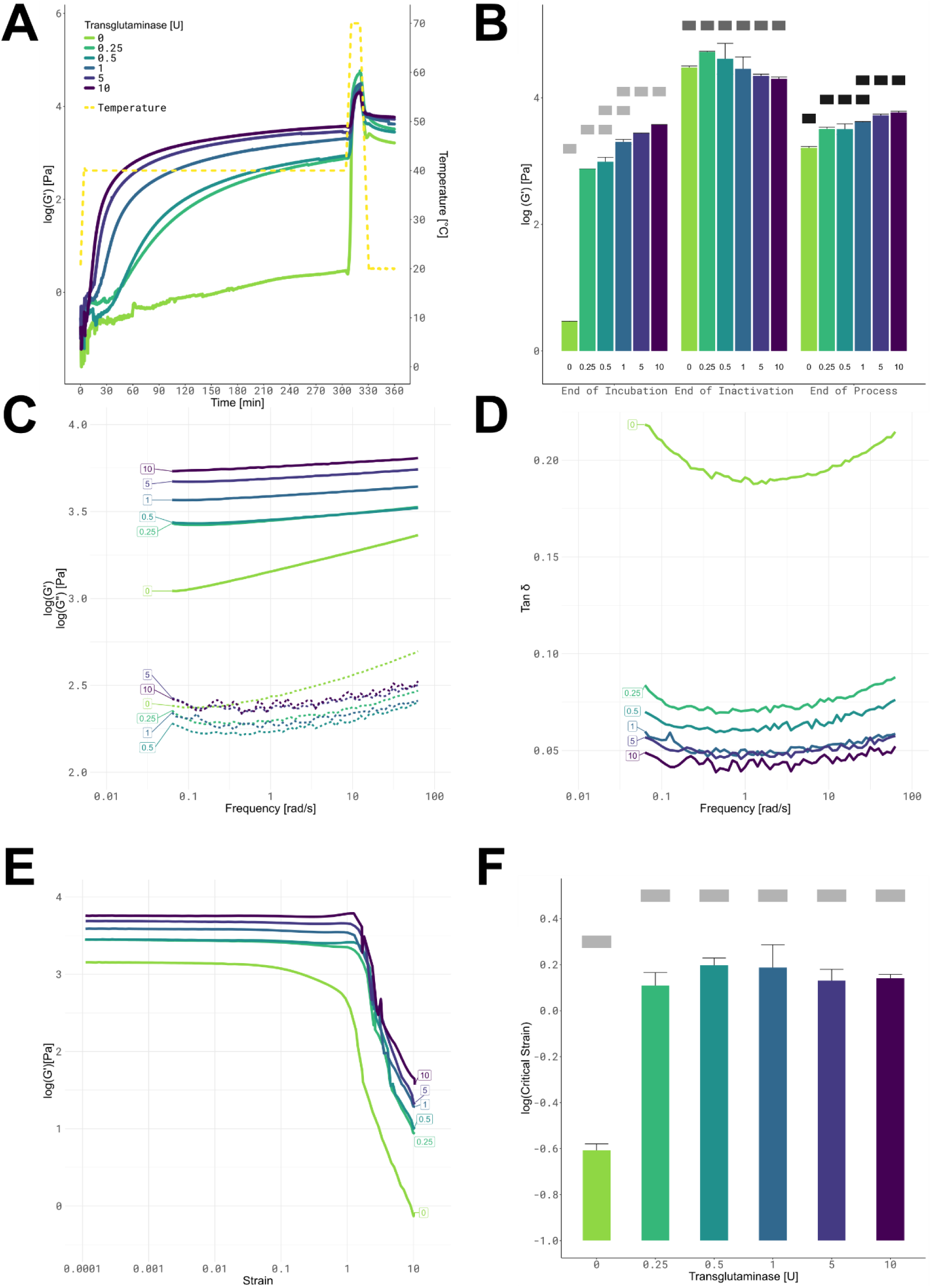
Rheological analysis of 10% (w/v) pea protein dispersions treated with 0, 0.25, 0.5, 1, 5, 10 U of TG/g protein at 40 °C and pH 7: **(A)** Time sweep showing the evolution of storage modulus (G′) over incubation and inactivation till process end. **(B)** G′ values during the time sweep at end of incubation, inactivation and process end. **(C)** Evolution of storage modulus (G′) (solid lines) and loss modulus (G”) (dashed lines) across frequency sweep **(D)** Tan δ (G″/G′) values across the frequency sweep. **(E)** Strain sweep depicting the linear viscoelastic region (LVR). **(F)** Critical strain for 0, 0.25, 0.5, 1, 5, 10 U TG/g protein. All G’,G” values shown are averages (measured in duplicate). The errors bars indicate standard deviation. Grey rectangular boxes at different levels indicate statistically significant differences (p<0.05).

G’ of the sample with 0 U TG (blank) showed limited change during incubation at 40 °C, whereas G’ of the samples incubated with TG increased significantly by the end of incubation (**Fig. 1 b**). It is noteworthy that, although the blank also shows gel formation due to the thermal inactivation process, the final G’ value of this system remained significantly lower (p< 0.05) than that of the samples incubated with the enzyme. Furthermore, it is also evident that the enzyme dosage had an effect on the gel properties of the pea protein. It follows from **Fig. 1 a** and **b**, that the rate of rise and the final value of G’ at the end of incubation and after cooling were higher with increasing enzyme dosages. Auer et al., (2025) reported an increase in G’ of commercial pea protein isolate due to TG crosslinking, but the effect was less noticeable for pea protein concentrate. However, they did not use a soluble native extract, so crosslinking might have been limited in their study. Furthermore, they also did not study a dose-response relationship between TG dosage and G’ evolution.

In addition to monitoring the rheological moduli of the systems during incubation and inactivation, the viscoelastic properties of the formed gels were evaluated using a frequency and strain sweep. A frequency sweep describes the mechanical spectra of a viscoelastic system within the LVR. The rheological moduli of the system were recorded over a broad frequency range where short and long time frames are represented by high and low frequencies, respectively (Moelants et al., 2014; Rao, 2014). **Fig. 1 c** depicts the frequency dependence of the G’ and G” of the protein gels with varying levels of TG added. Firstly, it was observed that the G’ value remains higher than the G” value across the frequency range for all the samples indicating retention of the gel structure in the system. The gels formed show characteristic behavior of covalently linked gels with little frequency dependence (Kavanagh & Ross-Murphy, 1998; Bayod et al., 2008). The frequency dependence of the G’ value could be quantitatively described by fitting a power law model (equation 1) into the data (Zhu et al., 2019). The R² (all > 0.97) values of the model fits showed that the power law accurately describes the rheological response of the protein gels (**Fig. 2 a** and **b**). The parameter estimates for the exponent (b) in the power law decreased, and that of the consistency factor (a) increased with increasing enzyme dosage, pointing towards decreasing frequency dependence and increasing gel strength (Ge et al., 2023). This indicated that increasing the dosage of TG reduces the frequency sensitivity of gels, thus forming a stronger network. Additionally, it could also be seen that the parameter estimates were significantly different for different enzyme dosages, confirming the importance of the enzyme dosage in developing structure in pea protein systems (**Fig. 2 a** and **b**).

**Figure 2.**
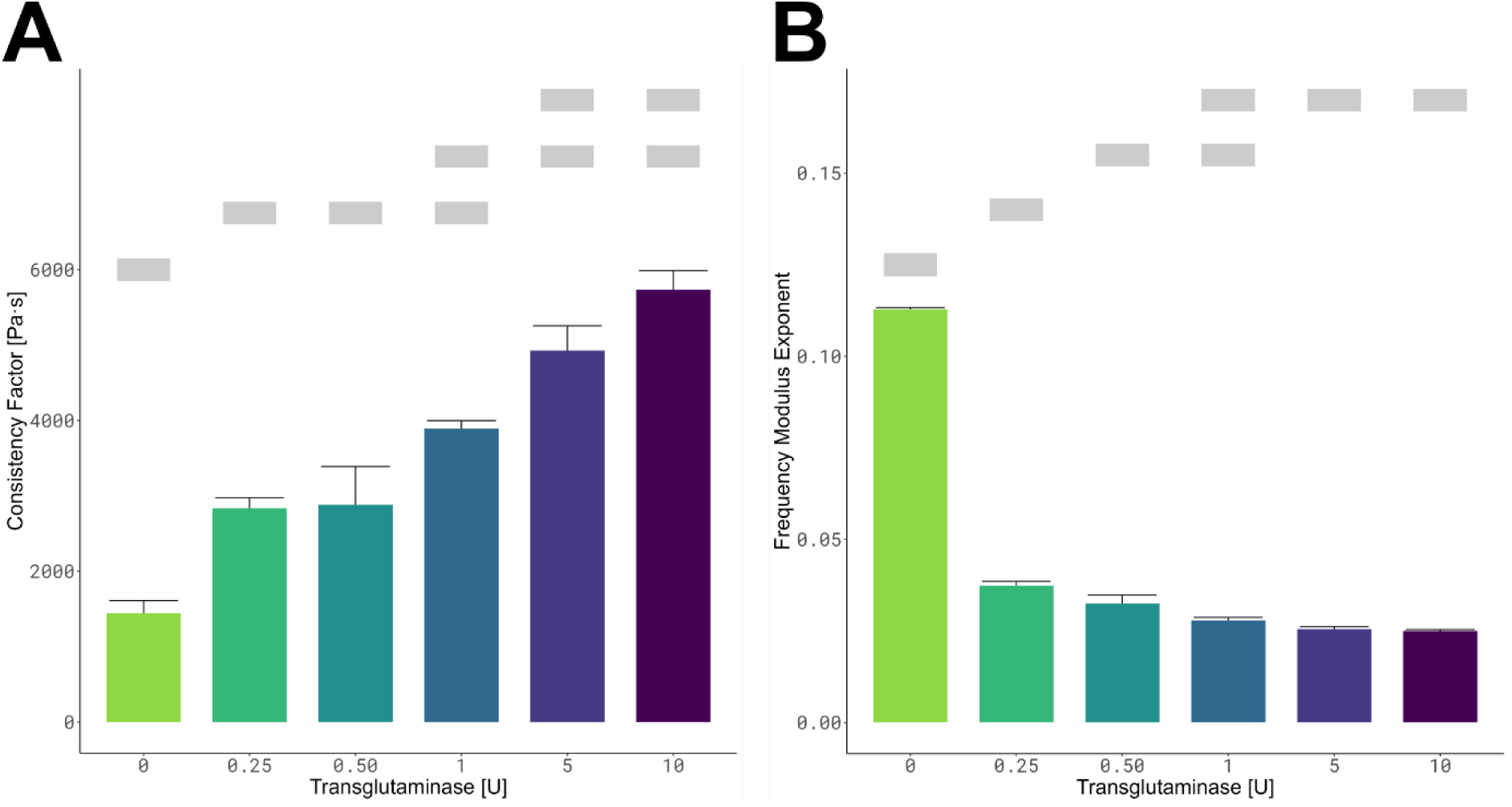
Frequency sweep parameters from the power law model (equation 1, section 2.5.4) of 10% (w/v) pea protein dispersions treated with 0, 0.25, 0.5, 1, 5, 10 U of TG/g protein at 40 °C and pH 7: **(A)** consistency factor (a, Pa-s^b^). **(B)** Frequency modulus exponent (b) All values shown are averages (measured in duplicate). The errors bars indicate standard deviation. Grey rectangular boxes at different levels indicate statistically significant differences (p<0.05).

Moreover, tan δ values (**Fig. 1 d**) are also important while interpreting frequency sweep data (Zhon et al., 2013). The tan δ values of the TG-treated samples remained rather constant across the frequency sweep. Additionally, it was also noted that the average tan δ remained higher than 0.1 for the blank and it decreased progressively with increasing enzyme dosage (0.2 ± 0.01 for the blank, 0.08 ± 0.01 for 0.25 U TG, 0.052 ± 0.003 for 1 U TG and 0.045 ± 0.003 for 10 U TG). The tan δ value of the crosslinked samples being lower than 0.1 across the frequency sweep is a characteristic of covalently linked strong gels (Kavanagh et al., 1998; Bayod et al., 2008).

Strain sweeps were also performed on the gel systems to determine the LVR. The LVR is defined as a range of strain values where G’ is independent of the imposed strain and is calculated as the strain value at which the G’ deviated by 10 % from the preceding five measurements (Glorieux et al., 2017). **Fig. 1 e** shows the behavior of the rheological moduli of the samples when subjected to a wide range of strain values. Increasing TG concentrations resulted in broadening of the LVR as evidenced by the critical strain values of the samples treated with different concentrations of TG. A similar observation was noted by Qin et al., (2022), who studied deformation behavior of pea protein treated with transglutaminase at conditions relevant for high moisture extrusion. The critical strain for the blank in the current study was 0.25 ± 0.02, whereas that of the samples treated with 0.25, 1 and 10 U TG were 1.3 ± 0.2, 1.6 ± 0.4 and 1.4 ± 0.1, respectively (**Fig. 1 f**). Moreover, it is also evident from **Fig. 1 e** that the irreversible deformation of the sample treated with 10 U TG occurred rather abruptly compared to the samples treated with lower TG concentrations and showed an upward trend at the end of the LVR. This is a typical characteristic of samples stabilized by a high density of covalent linkages and indicates the deformation resistance of the sample (Qin et al., 2022). Additionally, it was also observed that the G’ value in the LVR slightly increased with increasing enzyme dosage in the LVR as also seen at the end of the process in the time sweep (**Fig. 1 b**). To the best of our knowledge, this is the first time that an in-depth, comprehensive rheological characterization of pea protein gel properties as impacted by varying TG dosage has been reported.

### 3.3. Change in free primary amino groups due to crosslinking by TG

Since TG catalyzes acyl-transfer reactions between γ-carboxamide groups (acyl donor) of glutamine residues and free amino groups of lysine (acyl acceptor) in proteins, the concentration of free amino groups decreases with the progress of the crosslinking reaction. Thus, the change in concentration of primary amino groups can be interpreted as the progress of crosslinking, which can be quantified using the OPA method (Auer et al., 2025). The experimental data (Error! Reference source not found.) showed similar cross-linking behaviour across all enzyme dosages studied. The concentration of free amino groups decreased in the initial hour of incubation, followed by a receded rate of change. This suggests first-order kinetics, and the reaction curves were modelled using fractional conversion (equation 2) (Colle et al., 2010).

From the modelled kinetic data (Error! Reference source not found.), it could be observed that the estimated rate constant of the crosslinking reaction (*k*) increased with increasing enzyme dosage (0.007 min^-1^ ± 0.002 for 0.25 U TG/ g protein vs 0.017 ± 0.003 min^-1^ for 10 U TG/g protein).

**Table 1.** Parameter estimate of the rate constant, *k* (min^-1^) of pea protein crosslinking (10 % w/v) by transglutaminase (0, 0.25, 0.5, 1, 5, and 10 U transglutaminase/g protein) as a function of time estimated from the decrease in free amino groups due to the crosslinking reaction using the fractional conversion model (equation 1, section 2.6.2)

| TG dosage<br>(U/g protein) | $k$ (min<sup>-1</sup>) | | | | R<sup>2</sup> |
| --- | --- | --- | --- | --- | --- |
|  | Parameter<br>estimate | Standard<br>error | 95% Confidence<br>Limits |  |  |
| 0.25 | 0.007 | 0.002 | 0.002 | 0.012 | 0.82 |
| 0.5 | 0.007 | 0.001 | 0.005 | 0.009 | 0.94 |
| 1 | 0.009 | 0.002 | 0.005 | 0.014 | 0.93 |
| 5 | 0.014 | 0.001 | 0.012 | 0.015 | 0.99 |
| 10 | 0.017 | 0.003 | 0.011 | 0.022 | 0.93 |

In addition, the estimated final free amino group content also decreased progressively with increasing enzyme dosage, reaching 1.3 mg/g sample for 10 U TG, compared to 4.6 mg/g sample for 0.25 U TG after 300 min of incubation at 40 °C. This progressive decrease in free amino groups as function of TG concentration was also reported previously in pea protein, mung bean (Schlangen et al., 2023) and oat dough (Huang et al., 2010). These results are in line with the findings from the time sweep where the G’ of the samples increased faster and higher with increasing TG dosage at the end of incubation (section 3.2). The decrease of the free amino group content reached a plateau after 2 hours for the sample incubated with 10 U TG/g protein. For the lower enzyme dosages, the decrease of free amino groups slowed down, while continuing to progress until the end of the experimental time frame of 300 min. Possible reasons for this slowing down could be exhaustion of accessible substrate sites (for the higher enzyme dosages), or loss of enzyme activity over the experimental time frame (Schmidt et al., 1983; Robinson, 2015). A similar trend was also noticed by Djoullah et al., (2018), who observed a strong increase in the extent of crosslinking during the first two hours of the reaction. They also observed that denatured globulins showed higher extents of crosslinking compared to native ones. It should however be noted that the authors conducted the controlled denaturation was conducted in presence of salt (20 mM NaCl), which may have shielded electrostatic interactions, preventing proteins from aggregating too aggressively, thus retaining accessibility of the reaction sites to TG. Schäfer et al., 2007 reported poor correlation between decreasing free amino groups and rising ε-(γ-glutamyl) lysine isopeptide bonds, measured through high pressure liquid chromatography. This can be attributed to the fact that the OPA method is not specific to only detecting depletion of the free amino groups due to TG mediated crosslinking (Duijsens et al., 2023). Furthermore, masking of free amino groups within the gel network may result in inaccessibility of the OPA reagent to these residues (Flanagan et al., 2003). Therefore, more specific and in-depth analyses are required to fundamentally study the effect of TG mediated crosslinking reactions. To unravel participation of individual pea protein subclasses in the crosslinking reaction, in this study, SDS-PAGE analysis and proteomics approaches were used. These results are discussed in the subsequent sections.

### 3.4. SDS-PAGE analysis

The molecular mass profile of the protein extract was assessed using SDS-PAGE. As depicted in **Fig. 4**, the lanes corresponding to NHNE samples reveal that the most prominent bands observed were of 71 kDa (convicilin), 47 kDa, 17 kDa (vicilin), 40 kDa, 20 kDa (legumin bands) and less than 15 kDa (albumins) which conforms with literature expectations (Ladjal-Ettoumi et al., 2016). It is of note that the ultrafiltered protein studied in this work, consisted of a more complete protein profile including both globulins and albumins, as compared to extracts produced using a selective purification technique such as iso-electric precipitation which precipitates globulins only (R. Kornet et al., 2022) SDS-PAGE was also employed to monitor fading or disappearance of specific bands over the TG treatment time-course. This was based on the reasoning that these proteins form high-molecular-weight cross-link aggregates, which changes their molecular mass. The intensity of several protein bands decreased as a function of incubation time across all enzyme levels, indicating that the corresponding proteins were crosslinked and formed larger aggregates. It was further observed that certain protein bands appeared more susceptible to crosslinking, as shown by their more extensive disappearance with increasing incubation times. More specifically, bands of convicilin (71 kDa, vicilin (47 kDa), and acidic legumin (40 kDa) pea protein fractions were noticeably affected by TG treatment, whereas the alkaline legumin bands (20 kDa), and vicilin bands of 17 kDa showed limited intensity change, while albumins (less than 15 kDa) remained rather unaffected. Similar results were previously reported in faba beans, where this effect was attributed to differences in protein solubility (Nivala et al., 2017). Vicilins, for instance, are more hydrophilic than legumins, moreover, hydrophilic acidic-chains of legumin are located at the surface of the molecule legumin molecule while hydrophobic alkaline-chain sections are buried at the interior, restricting their solubilization in water and, as a result, the accessibility of TG (Karaca et al., 2011). Further evidence for protein cross-links is the appearance of new high molecular weight proteins bands at the very top of the SDS-PAGE gel in samples incubated with TG (**Fig. 4**). This stands in contrast to a lack of such high molecular weight bands in the NHNE and HNE samples, to which no TG was added. Thus, it can be inferred that the HMW protein aggregates were indeed formed due to TG-mediated crosslinking reactions.

**Figure 3:**
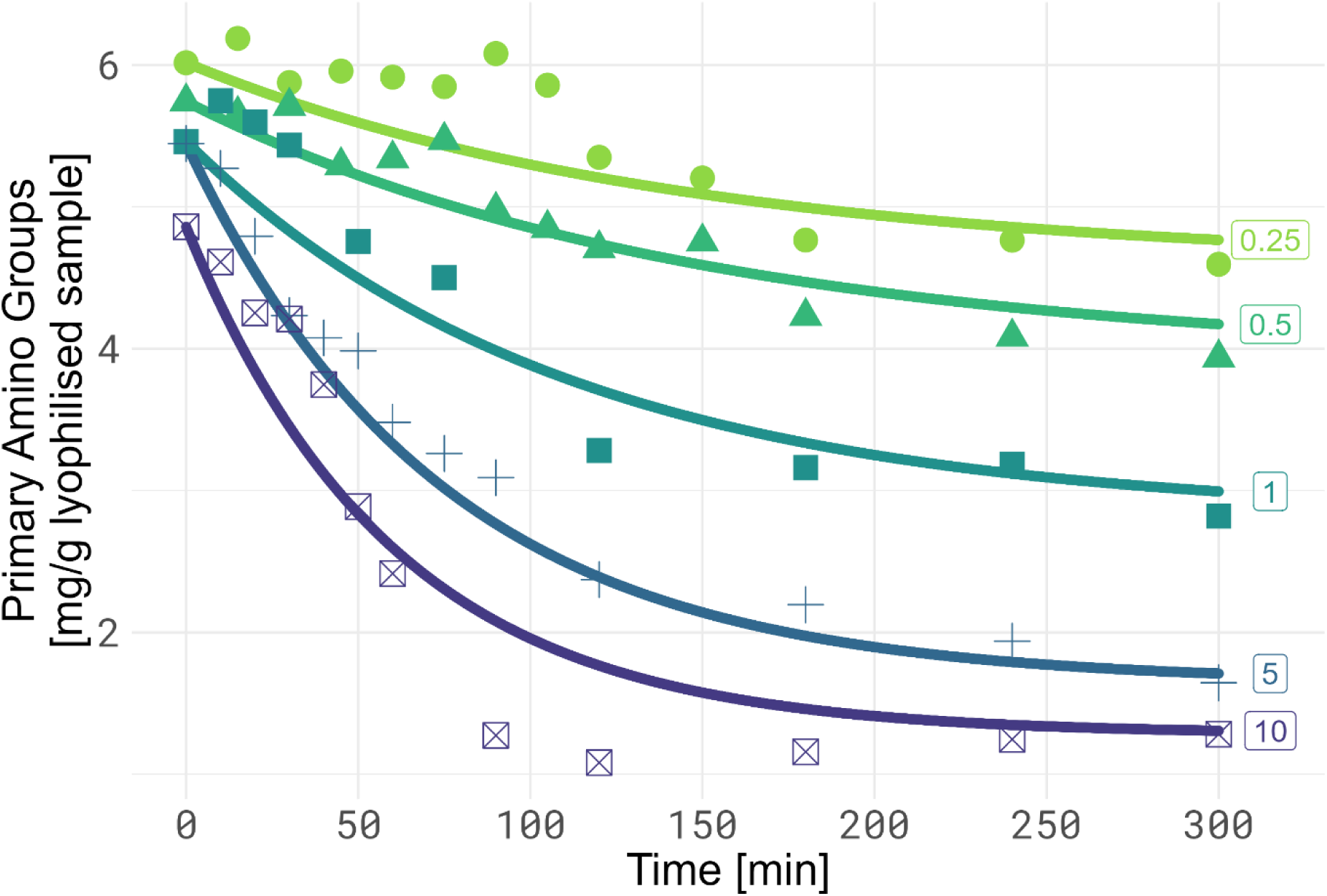
Change in primary amino groups in pea protein extract crosslinked by TG (0.25, 0.5, 1, 5, 10 U/g protein) at 40 °C, pH 7, as a function of incubation time. A fractional conversion model has been fitted to the kinetic data (equation 2, section 2.6.2). The symbols represent the experimental values and solid lines represent the modelled values.

**Figure 4.**
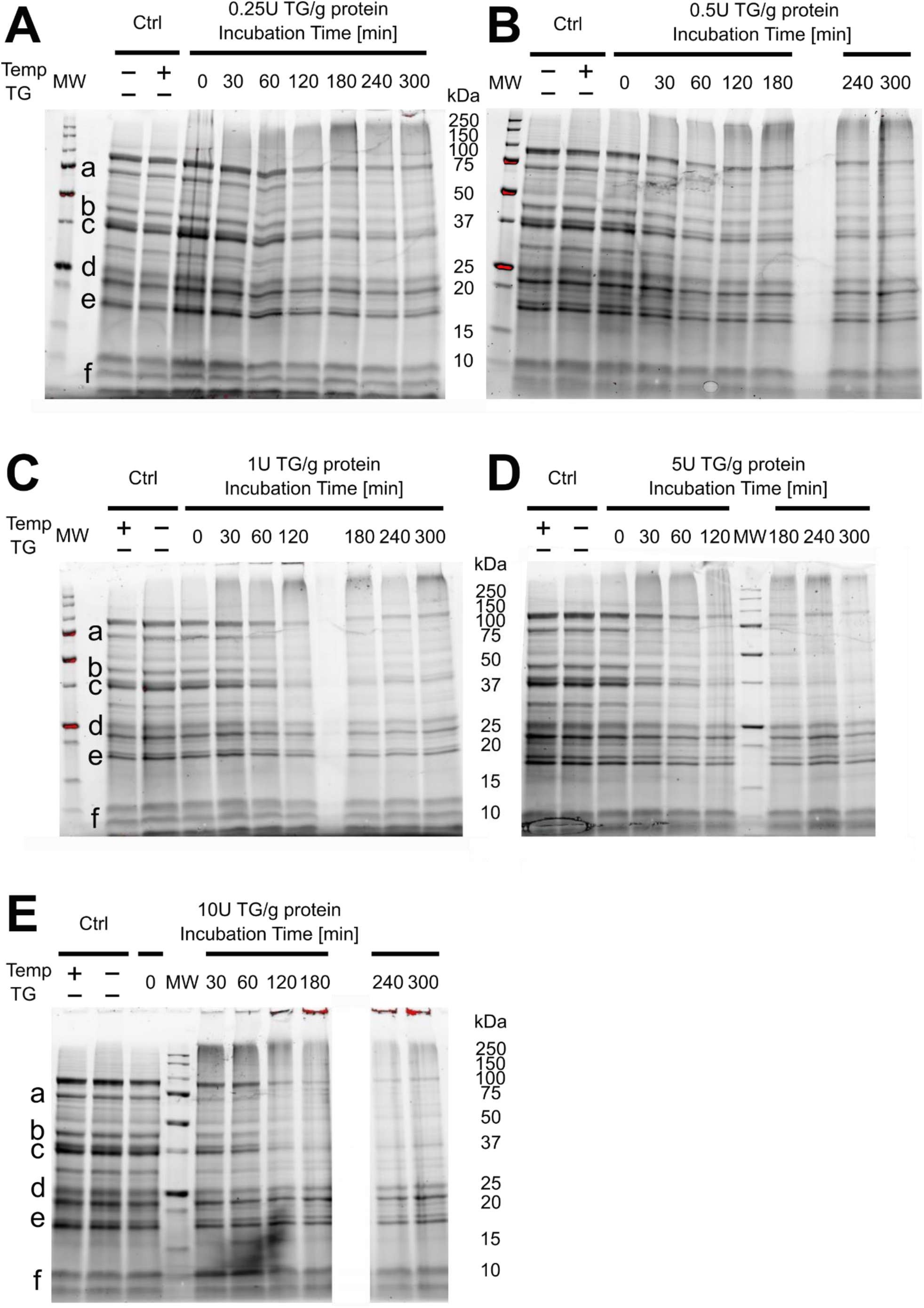
SDS-PAGE analysis of pea protein (10% w/v) treated with **(A)** 0.25, **(B)** 0.5, **(C)** 1, **(D)** 5 and **(E)** 10 U of TG/g protein at 40°C and pH 7, across various incubation times (0, 30, 60, 120, 180, 240, 360 min). Two control samples (Ctrl) were included: Lane 1 and 2 indicate whether heat was applied (Temp) and enzyme added (TG). Lanes 3-9 correspond to different incubation times. MW-Molecular weight markers.

It was also noticed that the band intensities of the participating protein subgroups at the end of the incubation time course were lower in the samples incubated with higher enzyme dosages, indicating higher levels of participation of the protein bands in the crosslinking reaction. Furthermore, the onset of disappearance of these protein bands was also earlier in the experimental time frame as the enzyme levels increased, indicating faster reaction rates. This is in accordance with the results from the rheological analysis and the OPA analysis, where, with increasing TG dosage, the G’ of the samples increase faster and higher, and the amino groups decreased faster and to a lower extent, respectively (section 3.2 and 3.3).

### 3.5. Mass spectrometric characterization of cross-links formed using proteome analysis

#### 3.5.1. Unfractionated cross-linked protein sample

Proteome analysis was used to gain more insight into the underlying molecular features of the cross-linked proteins, specifically an amino acid motif and protein domain features analysis was undertaken. Proteins from NHNE samples and samples after enzymatic crosslinking using 10 U TG/ g protein (n = 4) underwent proteome analysis. The first approach was to search the resulting MS data using pLink2, but this did not result in any cross-linked peptide identifications (data not shown). However, it was reasoned that participation of peptides in cross-linking leads to their lower abundance relative to their non-cross-linked precursors in the untreated samples. This means that peptides which are less abundant after TG treatment could be regarded as potentially cross-linked. Therefore, the MS data was re-searched using a non-cross-link-specialized search engine which resulted in the identification of 6799 unique peptides (≥ 95% probability) (**Supplement Table 1**). As seen in **Fig. 5 a**, an over-representation of lower-abundant peptides after TG treatment was detected, in accordance with the above stated hypothesis.

**Figure 5:**
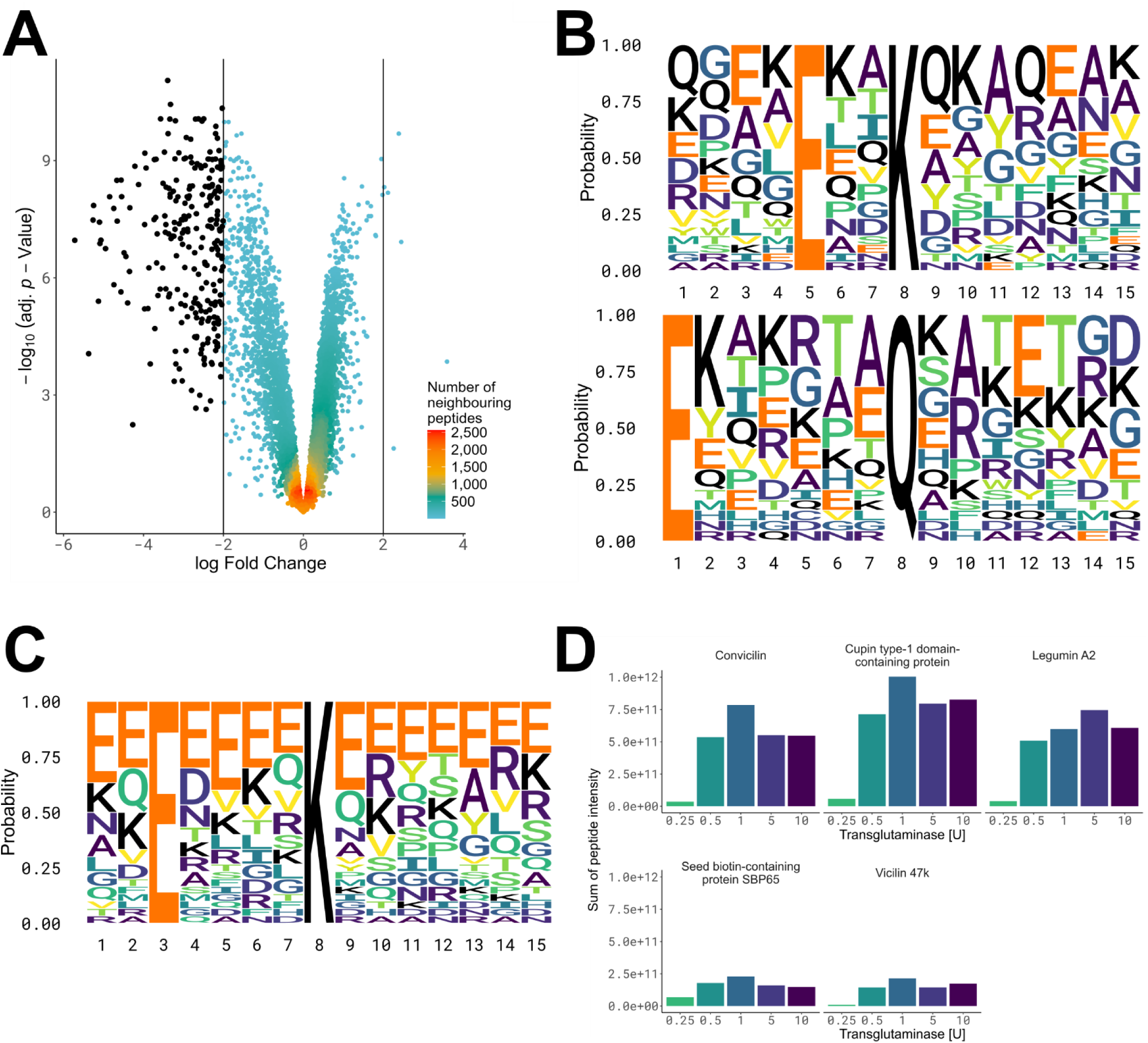
**(A)** Volcano plot depicting the change in peptide abundance between TG cross-linked and untreated control from whole protein lysate. Individual peptides are depicted. Black dots indicate over-representation of lower abundant peptides after TG treatment (negative fold change). **(B)** Motif analysis of the in black depicted peptides as seen in (A) with overrepresentation of glutamic acid E around K and Q sites. The MS raw files can be accessed from ProteomeXchange and jPOSTrepo under PXD064878 and JPST003865, while the analysis scripts to produce figure parts (A) and (B) can be accessed under https://doi.org/10.5281/zenodo.15640883. **(C)** Motif analysis of peptides exhibiting QR and/or QK adducts on lysine sites in the high molecular weight fraction (as per SDS-PAGE prefractionation) after TG crosslinking with validation of glutamic acid around potential K cross-linking site, as seen in (B). **(D)** Sum of MS1 intensities of peptides belonging to proteins with indicated names after treatment with TG at the indicated concentrations. Raw data for figure parts (C) and (D) can be accessed under PXD064875 and JPST003864, data analysis scripts used to produce these figure parts can be accessed via https://doi.org/10.5281/zenodo.15640016.

##### Identification of TG substrate motifs

The detected lower-abundant peptides were subjected to motif analysis to detect recurring amino acid sequence patterns surrounding TG targetable glutamine and lysine residues. Within the group of peptides potentially participating in cross-linking, 28 out of 94 exhibited a glutamic acid (E) at the third position before a potential lysine cross-link site (2.66-fold increase vs background, p < 2.66 x 10^-5^) and 22 out of 89 peptides had a glutamic acid (E) at the seventh position preceding glutamine cross-link site (2.76-fold increase vs background, p < 2.81 x 10^-5^), with further prominent appearance of glutamine in the close neighborhood surrounding the potential cross-link sites (**Fig. 5 b**).

##### Functional protein domain mapping

It is well known that sites of post-translational modifications lie in disordered regions of proteins, as these structurally flexible regions help the protein to better access the catalytic center of e.g., kinases (Bah et al., 2016). Similarly, it was also revealed that a substantial proportion of potential crosslinked sites were localized within a disordered region of the respective identified proteins. Potential cross-linked peptides containing a glutamine were positioned 4.50 times (p = 2.25 x 10^-21^) more frequently within disordered regions than within the set of all identified peptides (background), while for peptides containing lysine residues this was 5.34 times (p = 2.92 x 10^-15^). The potential importance of intrinsic disorder in substrate recognition for TG activity was also recognized before (Csosz et al., 2008). Previous studies reported that TG mainly induces protein crosslinking in the light meromyosin domain of the myosin rod (Feng et al., 2025). It was found that the flexible conformation of β-CN in casein micelles potentially enables better accessibility of substrate residues by TG (Hsieh & Pan, (2012). In line with these literature insights, the current study shows that flexible, disordered regions in which glutamine/lysine residues are structurally more accessible show a higher susceptibility to TG-mediated crosslinking.

#### 3.5.2. High molecular weight prefractionated cross-linked protein sample

The pLink2 search using the mass spectrometry data obtained from the unfractionated samples resulted in no identification of cross-linked peptide pairs (data not shown), and it was reasoned that this could be the result of the low frequency of cross-links within the peptide population. This can be overcome due to the fact that cross-linked proteins have an increased molecular mass and as such they can be enriched in the high molecular weight region of SDS-PAGE gels (**Fig. 4**). Two different strategies were applied for this. The data was searched using pLink2 (Chen et al., 2019) (**Supplement Table 2**) and further, a reporter mass-shift approach was undertaken. If a cross-linked glutamine (Q) is preceded or succeeded by a lysine (K) or arginine (R), this introduces an adduct on the cross-linked peptide containing the lysine cross-link site of either a deamidated QK (+257.1276 Da) or deamidated QR (+285.1437 Da), (**Supplement Fig. 1** and **Supplement Table 3**). There are no mono-isotopic mass adducts within the peptide search engine mass error window (20 ppm) reported in UniMod (Creasy & Cottrell, 2004), and they were thus deemed a suitable reporter mass-shifts for the identification of cross-links. A possible reporter mass-shift of +129.0790 Da could result from an adduct formed by a single deamidated lysine cross-linked to a glutamine. However, this mass is only 18 ppm smaller than that of a glycine (G) to tyrosine (Y) substitution (+129.0578 Da), which falls within the 20 ppm mass error allowed during the database search. For this reason, the deamidated K onto Q adduct was excluded. Using this approach, we identified 196 unique cross-linked peptides (5% FDR). The cross-linked amino acids were enriched within disordered protein regions (p = 4.76 x 10^-8^, 2.05-fold enrichment versus background) validating the results of the indirect search approach in section 3.5.1. However, a motif search did not result in any significant enrichments (data not shown). Via this approach, cross-links in vicilin (47 kDa), convicilin, and legumin A2 could be detected (**Supplement Table 4**), in line with their disappearance on the SDS-PAGE gels (**Fig. 4**). The reporter mass-shift approach resulted in 73 unique peptides for a QR remnant and 78 unique peptides for a QK remnant on lysine. Interestingly, although the search was against the complete pea protein FASTA containing 64176 entries, the majority of these mass adducts were placed on peptides belonging to proteins expected to be cross-linked, namely convicilin, legumin A2, and vicilin (47 kDa) (**Supplement Table 5)**. Further proteins with large numbers of peptides modified in such a way are cupin type-1 domain-containing protein (vicilin like protein) and Seed biotin-containing protein SBP65 (**Supplement Table 5)**. In total, 54 proteins exhibited reporter mass shifts, of which 20 overlapped with the 78 proteins detected via the cross-link search (**Supplement Table 6**). Out of these, 10 proteins (16 peptides) exhibited cross-link sites that were shared between the reporter mass-shift approach and the cross-link search (**Supplement Table 7**). Further, the amino acids exhibiting the reporter mass shifts were enriched in disordered regions (p = 5.05 x 10^-15^, 3.32 fold enrichment over background) and an enrichment of glutamic acid (E) at the -5 position relative to the modified lysine residue (p < 2.02 x 10^-5^, fold increase of 2.77 versus background), with further enrichment of E around the adduct site mimicking the results in section 3.5.1 (**Fig. 5 c**). This observation confirms a potential preference of transglutaminase for cross-linking within structurally flexible or disordered protein regions and enrichment of E around the substrate site favoring the formation of crosslinks, as discussed in section 3.5.1. Collectively, these findings highlight the reporter mass-shift approach as a valuable and biologically informative alternative to conventional cross-link search engines, capable of identifying TG cross-links while also providing insights into structural preferences of cross-linking sites. Next, in comparison with the OPA method, the relative amount of cross-linked peptides over time was established by quantitative mass spectrometry. Surprisingly, the quantification approach using TMT did not result in a time- and dose-dependent response for the number of detected crosslinks, as observed previously by OPA and rheological analyses (**Supplement Fig. 2**). MaxQuant and pLink2 only detect cross-linked peptide pairs. However, with prolonged incubation, it is likely that multiple peptides were cross-linked (with preference in the same disordered regions). This would lead to the same effect observed in the unfractionated cross-linked protein sample, i.e., the removal of these peptides from the identification pool with an effect on measured abundancy. This hypothesis would require that cross-linking sites could be established in the vicinity of each other. Indeed, several peptides exhibiting two reporter mass-shifts on the same peptide were identified (**Supplement Table 6** “Potential Multiple Cross-links”). Therefore, we applied an alternative approach, where it was reasoned that the total protein intensity (= sum of intensity of all measured peptides as per MS1 full scan) should be higher in samples treated with higher concentrations of TG, since a higher amount of cross-linking should result in more cross-linked protein in the HMW fraction. Using this approach, a higher intensity for convicilin, vicilin (47 kDa), and legumin A2 for 10 U TG was detected compared to 0.25 U TG. Further, as per the reporter mass-shift approach, Cupin type-1 domain-containing protein and Seed biotin-containing protein SBP65 also showed such a behavior (**Fig. 5 d**). These quantification results could potentially be explained by continued cross-linking by TG during the experimental time course. It could be noted that the bands that disappeared from the SDS PAGE during the course of crosslinking (section 3.4), also corresponded to the most abundant proteins in the crosslink search. However, proteomics enabled the identification of additional pea proteins that could not be identified using traditional analyses. Next, in comparison with the OPA method, the relative amount of cross-linked peptides over time was established by quantitative mass spectrometry. Surprisingly, the quantification approach using TMT did not result in a time- and dose-dependent response for the number of detected crosslinks, as observed previously by OPA and rheological analyses (**Supplement Fig. 2**). A possible explanation is that pLink2 only detects cross-linked peptide pairs. However, with prolonged incubation, it is likely that multiple peptides were cross-linked (with preference in the same disordered regions). This would lead to the same effect observed in the unfractionated cross-linked protein sample, i.e., the removal of these peptides from the identification pool with an effect on measured abundancy. This hypothesis would require that cross-linking sites could be established in the vicinity of each other. Indeed, several peptides exhibiting two reporter mass-shifts on the same peptide were identified (**Supplement Table 6** “Potential Multiple Cross-links”). Therefore, we applied an alternative approach, where it was reasoned that the total protein intensity (= sum of intensity of all measured peptides as per MS1 full scan) should be higher in samples treated with higher concentrations of TG, since a higher amount of cross-linking should result in the appearance of more cross-linked protein in the HMW fraction. Using this approach, a higher intensity for convicilin, vicilin (47 kDa), and legumin A2 for 0.5, 1,5 and 10 U TG was detected compared to 0.25 U TG. Further, Cupin type-1 domain-containing protein and Seed biotin-containing protein SBP65 also showed such a behavior (**Fig. 5 d**).

## 4. Conclusion

To the best of our knowledge, this is the first time that a comprehensive fundamental study reveals a macro- to molecular-level, insight into the structuring potential of pea protein using TG. The strength of the crosslinked gels increases significantly as a function of the TG dosage, thus directing the development of plant-based products with steered structural properties. Furthermore, while OPA analysis indicated a dose-response relationship between TG dosage and the level of crosslinking, SDS-PAGE, in agreement with the proteome analysis, further elucidated the responsiveness of convicilin, vicilin, and legumin subunits to the crosslinking reaction and the time-dependence of the reaction as well. Additionally, proteome analysis revealed preferential crosslinking of lysine and glutamine residues situated in the disordered regions of the protein and the presence of glutamic acid residues in the vicinity of the crosslink sites. Further, it was noted that the bands that disappeared from the SDS PAGE during the course of crosslinking (section 3.4), also corresponded to the most abundant proteins in the identified in the mass spectrometry crosslink search. Additionally, proteomics enabled the identification of additional pea proteins that could not be identified using traditional analyses. However, the quantification of cross-link formation is technically challenging due to the limitations of current search engines, where only cross-linked peptide pairs can be identified, while higher-order cross-links remain elusive. This influences the correct quantification of such peptides as cross-linking continues over time. A remedy would be to specifically hydrolyze all cross-links before mass spectrometry or during MS2. However, while such approaches are available for chemical cross-linkers, the usage of TG (as enzyme) renders this difficult. Based on the insights of this work, plant-based protein sources can be selected for structuring with TG. This selection can be done based on high contents of disordered regions with multiple lysine and glutamine residues, since based on the current study, these are potential substrates for TG-mediated crosslinking. Diversification in utilization of plant proteins is beneficial from an environmental and nutritional perspective. Moreover, the nutritional significance of the crosslinking reaction, i.e., the impact of TG-mediated crosslinking on protein digestibility and bio-accessibility, is also essential for further studies.

## Funding Acknowledgements

A. Mukherjee is a doctoral researcher funded by Flanders Innovation and Entrepreneurship (VLAIO) in the context of ProFuNu project (HBC.2021.0546). J. Van Pee is a doctoral researcher funded by Flanders Innovation and Entrepreneurship (VLAIO) in the context of ProFuNu project (HBC.2021.0546). D. Duijsens is a post-doctoral researcher funded by Research Foundation Flanders, Belgium (FWO - Grant no. 12A0225N).

## Acknowledgments

We would like to thank Steven Kasozi Galiwango for helping with the R script to extract TMT reporter intensities.

