## Supplement_Figures for "Transglutaminase mediated crosslinking of pea protein: from rheology to proteomics"

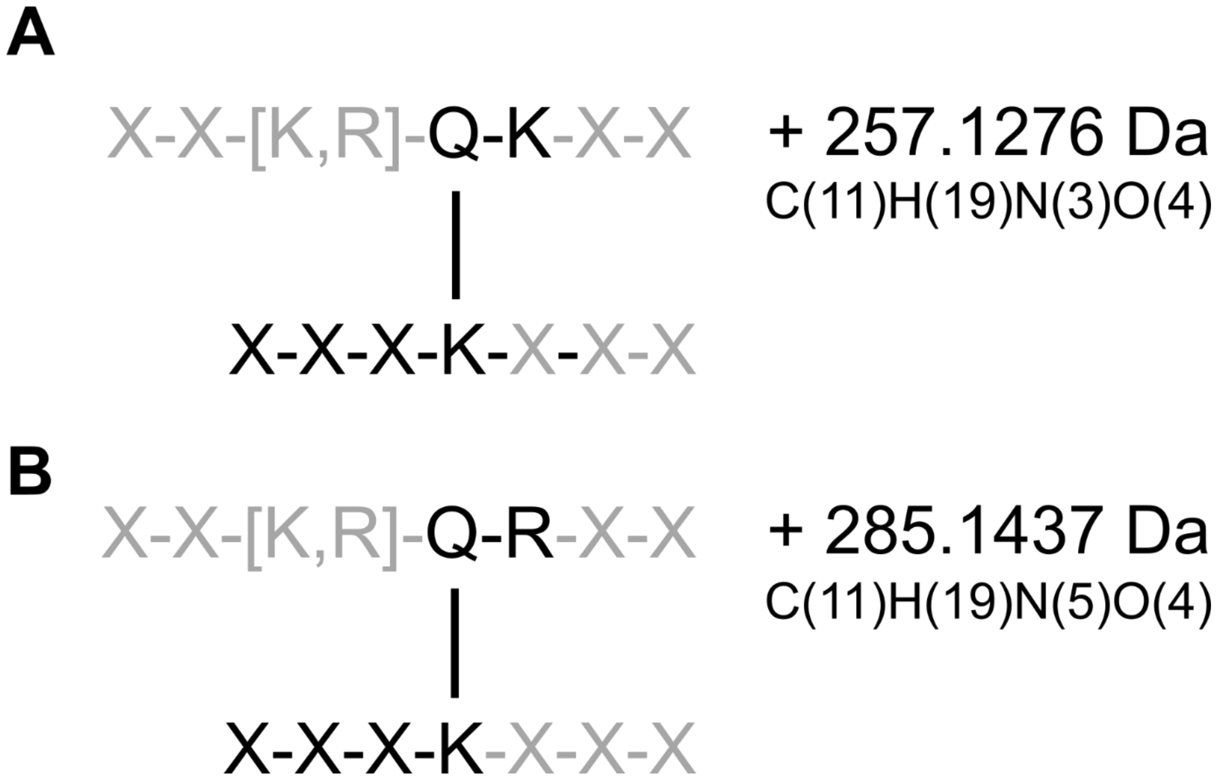


Supplementary Figure 1: Adduct formation on lysine for mass spectrometry reporter mass shift for the detection of potential cross-linking sites. **(A)** QK onto K and **(B)** QR onto K.

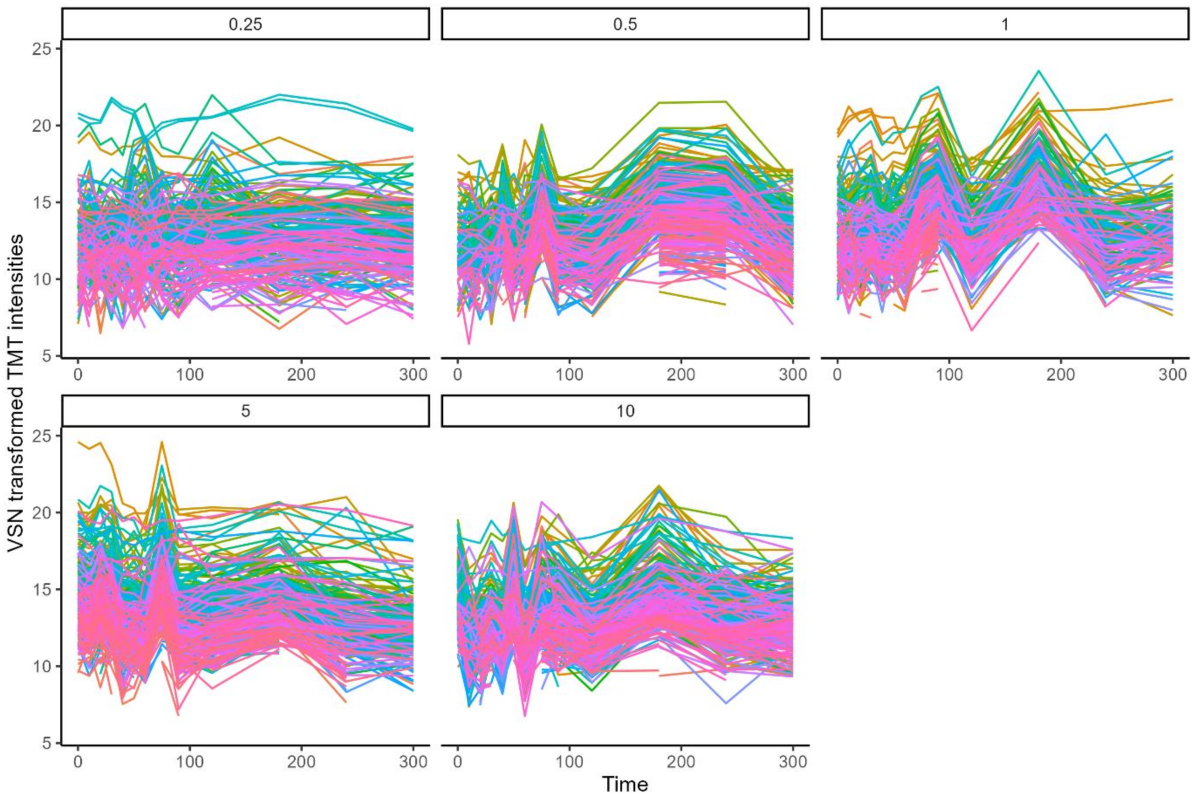


Supplementary Figure 2: TMT reporter intensities along the analysed time course. Number in figure titles depict the used TG concentration.
