## Supplement_Tables for "Transglutaminase mediated crosslinking of pea protein: from rheology to proteomics": Supplement_Table_Legends.docx

Supplement Table 01: iProphet output of Comet & TPP search of TG cross-linked whole protein lysate samples.

Supplement Table 2: pLink2 output of high molecular weight fraction (as per SDS-PAGE prefractionation) of TG cross-linked samples. Number after “DDA-” under “Title” indicates the MS run number, as per Table 2.

Supplement Table 3: MSFragger output of the adduct search (QK, QR) of the high molecular weight fraction (as per SDS-PAGE prefractionation) of TG cross-linked samples. Number after “DDA-” under “Title” indicates the MS run number, as per Table 2.

Supplement Table 4: Cross-linked peptides identified using pLink2 of the high molecular weight fraction (as per SDS-PAGE prefractionation) of TG cross-linked samples. MS run number as per Table 2, with indication of disordered regions

Supplement Table 5: MSFragger output of the adduct search (QK, QR) of the high molecular weight fraction (as per SDS-PAGE prefractionation) of TG cross-linked samples. MS run number as per Table 2, with indication of disordered regions.

Supplement Table 6: Overlap between pLink2 and MSFragger adduct search of high molecular weight fraction (as per SDS-PAGE prefractionation) of TG cross-linked samples, indicated as “Overlap.Cross.Link.and.Adduct” = TRUE. Leading Number separated by “.” under “Scan.Number” indicates the MS run number as per Table 2.

Supplement Table 7: Overlap between pLink2 and MSFragger adduct search of high molecular weight fraction (as per SDS-PAGE prefractionation) of TG cross-linked samples. Leading Number separated by “.” under “Scan.Number” indicates the MS run number as per Table 2.
